# Insight into the scaffolding function of USP18 from a high resolution cryo-EM structure of STAT2-USP18-ISG15 ternary complex

**DOI:** 10.64898/2026.02.12.705587

**Authors:** Kevin W. Huynh, Rachel Plumb, David R. Healy, Veronica Jové, Erik C. Ralph, Chiachin Wilson Lee, Heather Wheeler, Kymberly Levine, Zhen Huang, Kimberly Fennell, Richard A. Corpina, Timothy Craig, Paul D. Wes, Paula M Loria, Monica Schenone, Seungil Han, Feng Wang, Huixian Wu, Masaya Yamaguchi

## Abstract

USP18 is a primary negative regulator of the type I interferon (IFN-I) signaling which regulates hundreds of IFN-stimulated genes for viral protection and anti-cancer immunity. USP18 plays dual roles in the IFN-I signaling: 1) deubiquitinase enzymatic function which cleaves ISG15 from its substrates and 2) scaffolding function through forming a complex with STAT2 to suppress IFN-I signaling. Targeting the scaffolding function of USP18, instead of its enzyme activity, is crucial for reducing cancer cell fitness and boosting anti-tumor immunity. However, the molecular basis of USP18’s scaffolding function remains unclear due to the lack of structural information. Here, using a fusion tag strategy, we captured the transient USP18-STAT2 complex and determined a ternary complex structure of STAT2-USP18-ISG15 at 3.05 Å resolution by cryogenic electron microscopy (cryo-EM) that delineated detailed USP18-STAT2 interactions. Remarkably, the ternary complex impairs USP18’s enzymatic function by STAT2-mediated disruption of its catalytic triad. Structural analysis and mutagenesis identify specific USP18 point mutations, facilitating further investigation into the role of USP18 in IFN-I signaling. Taken together, our findings suggest that USP18’s scaffolding function could present an untapped opportunity for cancer therapy.

## Introduction

Type-I interferon (IFN-I) signaling is an indispensable defense mechanism for viral or pathogen infections and is also essential regulators for cellular immunity^1–6^. Upon IFN binding to the surface receptors IFNAR1 and IFNAR2, the receptor-associated protein tyrosine kinases Janus kinase 1 (JAK1) and tyrosine kinase 2 (TYK2) phosphorylate signal transducer and activator of transcription 1 and 2 (STAT1/STAT2), resulting in a heterotrimeric complex with Interferon regulatory factor 9 (IRF9), an IFN-stimulated gene factor 3 (ISGF3) complex. The ISGF3 complex translocates into the nucleus and induces hundreds of IFN-stimulated genes (ISGs) by engaging IFN-stimulated response elements in ISG gene promoters. Once expressed, ISGs play a central role in antiviral protection. Besides the IFN-induced antiviral protection, IFN-I signaling also plays a pivotal role in antitumor immunity by enhancing anti-cancer immune responses which include boosting antigen-presentation, stimulating immune cells and eliminating tumor cells^3–5^. Combining the PD1/PD-L1 blockade with IFN also enhances anti-cancer immunity^7,8^. Therefore, modulating key regulators of IFN-I signaling may offer an effective approach to prevent cancer cell growth.

Upon activation of IFN-I signaling, interferon-stimulated gene 15 (ISG15) is conjugated to various substrates via a mechanism analogous to the ubiquitin (UB) E1-E2-E3 cascade (UBA7-UBCH8-HERC5). This mechanism is known as ISGylation^9–13^. Ubiquitin-specific peptidase 18 (USP18) has been identified and characterized as a deubiquitinase enzyme, but it specifically removes ISG15 from various ISGylated substrates, a process known as deISGylation^14–17^. Recent genetical and biochemical studies have identified an additional role for USP18 in the IFN-I signaling^18,19^. Specifically, USP18 acts as a scaffold that blocks downstream JAK/STAT signal activation through direct interactions with STAT2 and IFNAR2^20,21^. USP18 is recruited to IFNAR2 via interactions with STAT2 coiled-coil (CC) and DNA-binding domains; disrupting this USP18-STAT2 interaction increases sensitivity to IFN stimulation^20^. Point mutations in USP18 (I60N) and STAT2 (R148W or R148Q) derived from type I interferonopathy patients which disrupt USP18-STAT2 interactions were shown to cause prolonged JAK-STAT signaling and transcriptional activations^22–24^. Our earlier analysis found that USP18’s scaffolding function has a greater effect than its catalytic function in regulating IFN-I mediated immune responses, suggesting possible therapeutic potentials^25^. Nevertheless, the absence of structural data regarding USP18-STAT2 interactions constrains our understanding of USP18’s scaffolding role in negative regulation of IFN-I induced JAK/STAT activation.

Here, we elucidated the structural mechanisms underlying the scaffolding function of USP18 in the IFN-I signaling by solving the cryo-EM structure of the human STAT2-USP18-ISG15 ternary complex. The experimental ternary complex structure reveals key USP18-STAT2 interactions which were not predicted in the AlphaFold2 model. The ternary complex structure and biochemical characterization reveal how STAT2 regulates catalytic activity of USP18 at the molecular level. These findings define the distinct functions of USP18 and may aid drug discovery targeting USP18 and future research on its function in IFN-I signaling.

## Results

### STAT2-USP18-ISG15 ternary complex structure by cryo-EM

Our previous work demonstrated that disrupting the scaffolding function of USP18 restored IFN sensitivity in human cancer cells, while its catalytic function did not, as shown by comparing the patient-derived USP18 I60N mutant with the catalytically inactive USP18 C64S mutant^25^. It underscored the crucial role of USP18’s scaffolding function in cancer cell proliferation. To investigate the interaction of USP18 with STAT2 and ISG15 in its scaffolding role, we used cryo-EM to resolve the wild-type (WT) STAT2-USP18-ISG15 ternary complex. This effort, however, only resulted in a low-resolution reconstruction, suggesting flexibility or transient interactions between USP18 and STAT2 (Fig. 1a, Supplementary Fig. S1a). This transient nature was further supported by mass photometry revealing both ternary complex and dissociated populations in the WT complex when the proteins were co-expressed in insect cells (Fig. 1b).

**Fig. 1.**
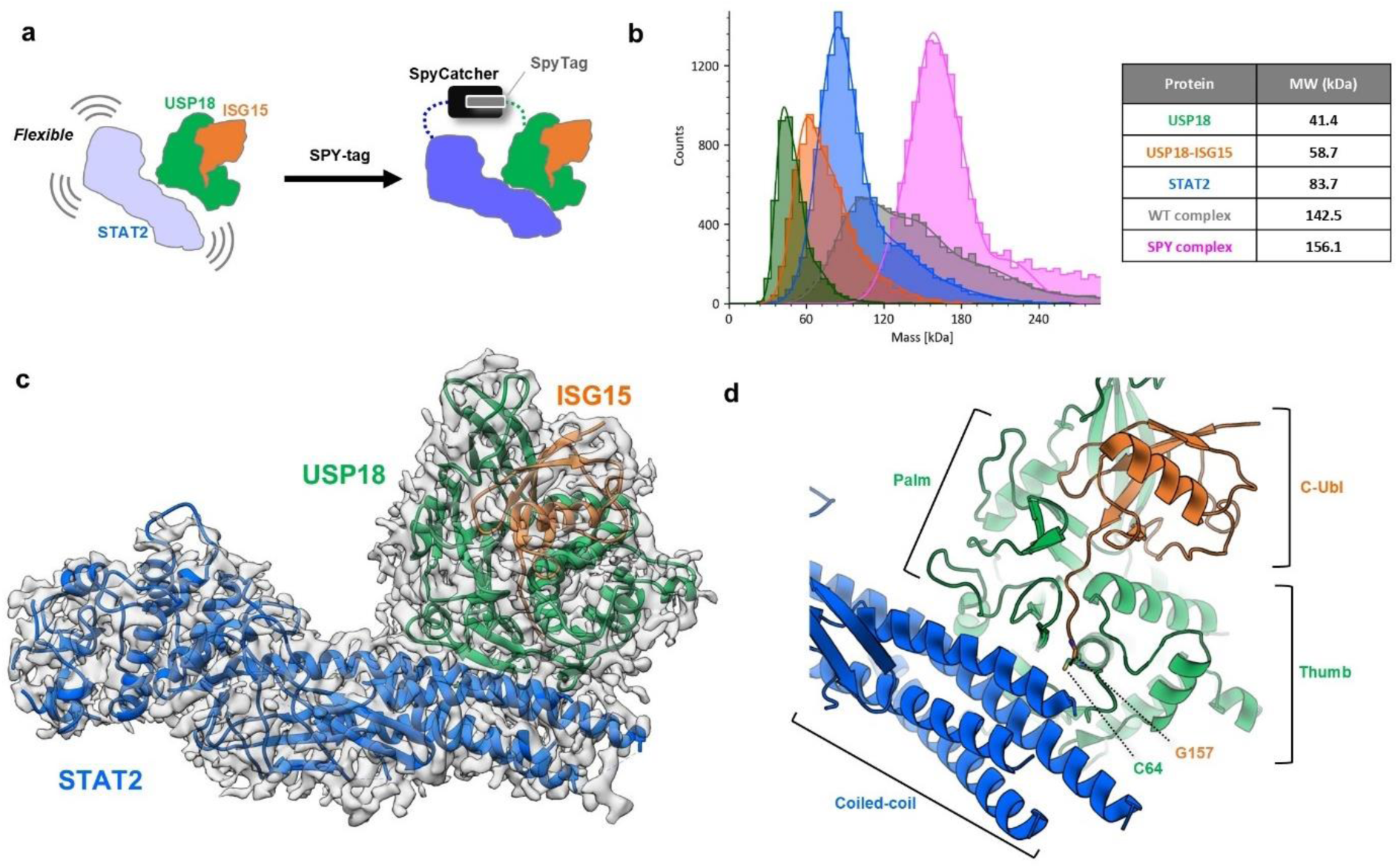
Structural determination of STAT2-USP18-ISG15 complex by cryo-EM. **a**) The scheme to stabilize the STAT2-USP18-ISG15 ternary complex by SPY-tag. **b**) Mass photometry characterization of cryo-EM samples. Theoretical molecular weight is shown in the table. **c)** The overview of STAT2-USP18-ISG15 complex structure with cryo-EM maps. STAT2, USP18 and ISG15 are colored in blue, green, orange respectively. **d)** Close up view of domains or regions involved in the USP18-STAT2 interactions. USP18 C64 and ISG15 G157 are shown with a stick model.

Given the strong affinity between USP18 and ISG15^25^, we attributed the observed dynamics mainly to the weak interaction between USP18 and STAT2, presenting a tremendous challenge to enable a high-resolution cryo-EM structure of the intact ternary complex using the native system. To stabilize the transient USP18-STAT2 interactions, we applied a covalent fusion tag, SpyCatcher/SpyTag (SPY-tag) that spontaneously couples, when in close vicinity, a 13 amino acid peptide SpyTag with a 11 kDa SpyCatcher protein originally derived from FbaB in *Streptococcus Pyogenes*^26,27^. We leveraged the STAT2-USP18-ISG15 complex model previously generated by AlphaFold2^25^ (AF2 model) to design SpyCatcher at the N-terminus of STAT2 and SpyTag at the N-terminus of USP18 (Supplementary Fig. S1b, S3a). The isopeptide bond formation between SpyCatcher-STAT2 and SpyTag-USP18 was confirmed by sodium dodecyl sulfate polyacrylamide gel electrophoresis (SDS-PAGE) (Supplementary Fig. S1a). Furthermore, the SPY-tagged USP18-STAT2 co-expressed with ISG15 can be purified into a homogeneous ternary complex as demonstrated by the single peak in mass photometry (Fig. 1b).

The SPY-tag facilitated stabilization of the ternary complex, enabling determination of the STAT2-USP18-ISG15 ternary complex structure by cryo-EM at a resolution of 3.05 Å (Fig. 1c, Supplementary Fig. S2a). Most of the USP18 finger, palm and thumb regions were resolved showing that USP18 wraps around ISG15 to form a fist-like structure. The coiled-coil (CC) domain, DNA-binding domain (DBD), linker-domain (LD) and SRC homology 2 (SH2) domains of STAT2 were also resolved by the cryo-EM densities resembling the forearm part of the structure (Fig. 1c-d, Supplementary Fig. S2b). The C-terminal ubiquitin-like (C-Ubl) domain of ISG15 was unambiguously modeled into the cryo-EM densities, but the N-terminal Ubl (N-Ubl) domain was not resolved, suggesting flexibility of this domain likely due to the lack of interaction with USP18 in the complex (Supplementary Fig. S2b, S3b). This is consistent with the mouse USP18-ISG15 complex structure, where mISG15 N-Ubl showed minimal interaction with mUSP18^28^. The cryo-EM structure showed that the palm/thumb regions and the catalytic module of USP18 interact mainly with the STAT2 CC domain near “the wrist” of the structure, covering about 2900 Å^2^ of solvent-accessible surface area (Fig. 1c-d, Supplementary Fig. S2b). By contrast, ISG15 demonstrates minimal interaction with STAT2, with the only contact observed made by the C-terminal G157 buried between the palm and thumb of USP18 and STAT2 CC domain (Fig. 1d).

### Details of USP18-STAT2 interactions

While the overall structural arrangement for USP18-ISG15 binary complex in the cryo-EM structure largely agrees with the AF2 model, the USP18 and STAT2 conformations deviate from the observed AF2 model (Supplementary Fig. S3). The root-mean-square-deviation (RMSD) between the cryo-EM structure and the AF2 model for the entire ternary complex and the USP18-ISG15 binary complex is 2.6 Å and 0.8 Å respectively. The key structural differences are observed at the STAT2 CC domain (shifted 2-4 Å) and the STAT2 LD-SH2 domain (shifted ∼10 Å) between the cryo-EM structure and the predicted model (Supplementary Fig. S3c-d). As a result, the cryo-EM structure uncovered new interactions mediated by side chains of STAT2 and USP18 that were not predicted by the AF2 model.

There are various reported physiological USP18 and STAT2 mutations that disrupt the USP18-STAT2 complex and result in IFN-I-mediated autoinflammation and type I interferonopathy^22–25,29^. The STAT2 mutations R148W and R148Q were found in patients with type I interferonopathy^23,24^. In the cryo-EM structure, STAT2 R148 forms a salt bridge with D346 and hydrogen bonds with S343 in the palm region of USP18. Furthermore, this residue facilitates a large polar interaction network involving USP18 Y322 and STAT2 R223 and T226 (Fig. 2a-b, Supplementary Fig. S2c). Mutant STAT2 R148Q likely disrupts the salt bridge with D346 of USP18 due to the replacement of a positively charged residue to an uncharged residue, while the bulky side chain of R148W causes steric hinderance. Both mutations lead to USP18 displacement from STAT2 which results in releasing the negative regulation of USP18 on the IFN-I signaling^23,24^. USP18 I60N was previously characterized as a scaffolding mutant that mimics the genotype of a USP18 knock-out (KO)^22,25^. USP18 I60 is located on a hydrophobic patch of the thumb region near the active site, interacting with the hydrophobic cluster L203, L206 and P292 of STAT2 CC domain (Fig. 2c, Supplementary Fig. S2d). The I60N mutation perturbs these hydrophobic interactions between STAT2 and USP18 owing to the substitution of a hydrophobic residue to a polar residue. STAT2 A219V mutant was also reported to disrupt USP18 interactions^29^. Based on the cryo-EM structure, mutating STAT2 A219 to Val likely cause a steric clash with USP18 N338, leading to disruption of the USP18-STAT2 interaction (Fig. 2d, Supplementary Fig. S2e).

**Fig. 2.**
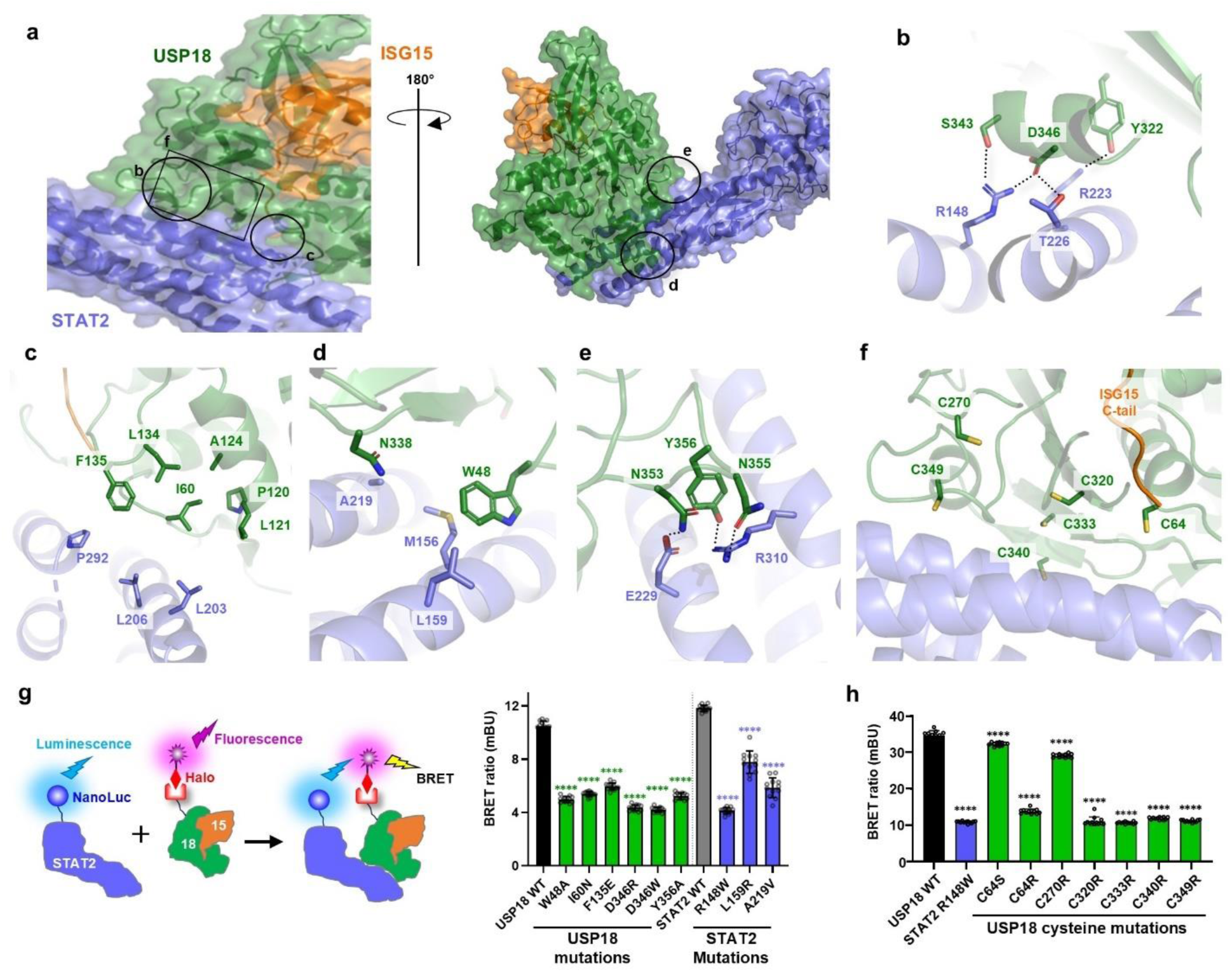
Details of USP18-STAT2 interactions. **a)** Selection of unique USP18-STAT2 interactions in the cryo-EM structure (b-f). **b)** STAT2 R148, T226 and R223 electrostatic interactions with Y322, S343, D346. **c)** USP18 I60 hydrophobic patch interactions with STAT2 CC domain harboring L203, L206, P292. **d)** USP18 W48 hydrophobic interactions with STAT2 M156 and L159. STAT2 A219 is located near USP18 N338. **e)** The electrostatic interaction between USP18 N353/N355/Y356 and STAT2 E229/R310. **f)** USP18 surface cysteine patch (C64, C270, C320, C333, C340, C349) near the STAT2 interface. **g)** NanoBRET assay of the residues involved in the USP18-STAT2 interface shown in b-e. The scheme of assay is shown left. Data are mean ± s.d. (n = 12, biologically independent samples) with P-values obtained by ordinary one-way ANOVA test; green **** indicate significance at P < 0.0001 to USP18 WT, blue **** indicate significance at P < 0.0001 to STAT2 WT. **h)** The NanoBRET assay for USP18 cysteine patch shown in f. Data are mean ± s.d. (n = 12, biologically independent samples) with P-values obtained by ordinary one-way ANOVA test; **** indicate significance at P < 0.0001.

The cryo-EM structure not only explained how reported physiological mutations affect the USP18-STAT2 interaction and the scaffolding role of USP18, but also identified additional residues critical for this complex formation. To understand the effects of these residues on the USP18-STAT2 interface, we established a cellular NanoBRET (Nano-Luciferase Bioluminescence Resonance Energy Transfer) assay where NanoLuc-tagged STAT2 transfers bioluminescence energy to the fluorescently labeled Halo-tag ligand on Halo-tagged USP18 when the complex forms (Fig. 2g). We confirmed by the NanoBRET assay that the STAT2 R148W mutant showed a significant reduction in BRET ratio consistent with a disruption of the USP18-STAT2 complex formation^24^. Additionally, USP18 D346W and D346R mutants showed a similar decrease in the BRET ratio as STAT2 R148W, supporting its essential role in complex formation via interactions with STAT2 residues R148 and T226 as observed in the structure (Fig. 2g). A hydrophobic pocket was observed near the STAT2 A219V mutant consisting of residues USP18 W48 and STAT2 M156 and L159 (Fig 2d, Supplementary Fig. S2e). Mutations introduced in this hydrophobic pocket such as USP18 W48A and STAT2 L159R showed a reduction in BRET ratio, supporting the key role of these hydrophobic interactions in USP18-STAT2 complex formation (Fig. 2g). Similarly, the USP18 I60N and F135E substitutions which affect the hydrophobic interaction with STAT2 hydrophobic patch containing L203, L206 and P292 showed a marked decrease in BRET ratio (Fig. 2g). Extensive electrostatic interactions are also observed between charged residues lining the USP18-STAT2 interface. Specifically, USP18 residues N353, N355 and Y356 formed ionic interactions with STAT2 residues E229 and R310 (Fig. 2e, Supplementary Fig. S2f). The mutation Y356A of USP18, which abolished interactions with STAT2 R310 led to a significant reduction of USP18-STAT2 NanoBRET signal, confirming the important role of the Y356-R310 interaction in the complex formation (Fig. 2g).

A patch of surface cysteine residues in USP18 (C64, C270, C320, C333, C340 and C349) lines the USP18-STAT2 interface (Fig. 2f). Single point mutations of these cysteine residues to bulky, positively charged arginine disrupts the electrostatic interactions between negatively charged USP18 and positively charged STAT2 (Supplementary Fig. S2g), leading to a significant decrease in the BRET ratio, except for C270 (Fig. 2g). We reasoned that mutating USP18 C270 did not affect the interaction as the residue is too far away from the USP18-STAT2 interface (Fig. 2f, 2h, Supplementary Fig. S2g). In addition, unlike C64R, mutating C64 to a small, uncharged Ser residue C64S did not significantly reduce the BRET ratio, suggesting USP18 bearing Ser at the catalytic C64 can still bind to STAT2 (Fig. 2h). In summary, our cryo-EM analysis of STAT2-USP18-ISG15 complex and mutagenesis experiments reveal pivotal USP18-STAT2 interactions playing a role in the USP18’s scaffolding function. The structure also clarifies the impact of previously reported physiological mutations in USP18 and STAT2 in the context of the complex formation.

### Role of ISG15 in USP18-STAT2 interactions

It has been reported that the USP18-ISG15 interaction is necessary, but not sufficient for controlling IFN-I signaling as shown in studies using ISG15 C-terminal Gly-Gly mutants, which do not bind to USP18^30^. Thus, we also sought to understand whether the USP18-ISG15 interactions are vital for the USP18-STAT2 interactions using the NanoBRET assay. For this, we leveraged a USP18 mutant bearing the mutations (A141Q/L145R/P195Q/H255Q) in its ISG15 binding box 1 (IBB1), which was reported to weaken the USP18-ISG15 interaction (Fig. 3a)^28^.

**Fig. 3.**
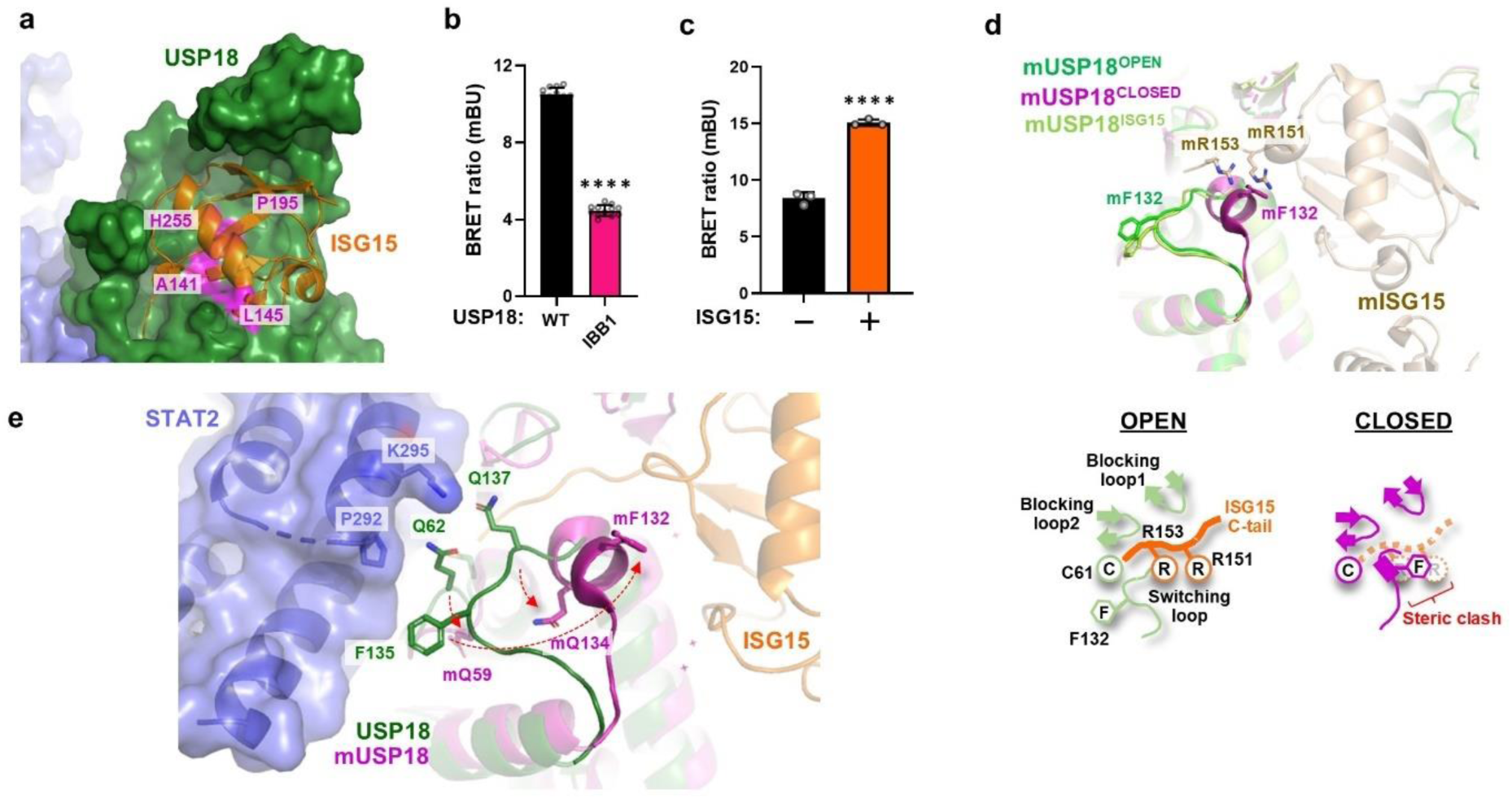
Role of ISG15 in USP18-STAT2 interactions. **a)** Location of ISG15-binding box 1 (IBB1: A141Q/L145R/P195Q/H255Q) (magenta) in the cryo-EM structure. **b)** NanoBRET assay of USP18 IBB1 mutant. Data are mean ± s.d. (n = 12, biologically independent samples) with P-values obtained by ordinary one-way ANOVA test; **** indicate significance at P < 0.0001. **c)** NanoBRET assay either in the absence (-) or presence of 1 μg (+) ISG15. Data are mean ± s.d. (n = 3, biologically independent samples) with P-values obtained by unpaired t-test; **** indicate significance at P < 0.001. **d)** Distinct switching loop conformations in USP18 OPEN and CLOSED states. Two protomers of apo mUSP18 (PDB 5CHT) and mISG15 bound mUSP18 (PDB 5CHV) are overlayed. Apo mUSP18 in OPEN and in CLOSED states are colored green and magenta while mUSP18 in a complex with mISG15 (OPEN) is colored in lime. Cartoon representation for OPEN/CLOSED states are depicted below. **e)** The conformational diversity of USP18 OPEN and CLOSED states mapped on the cryo-EM structure. CLOSED state of mUSP18 derived from apo mUSP18 (PDB 5CHT) (magenta) was overlaid.

The USP18 IBB1 mutant, like other USP18 mutants, reduced the NanoBRET signal, indicating that the USP18-ISG15 interaction enhances USP18-STAT2 affinity (Fig. 3b). Additionally, the NanoBRET result showed that the USP18-STAT2 interactions were markedly reduced in the absence of ISG15 (Fig. 3c). Structural comparison between the cryo-EM structure and previously reported mUSP18 structures (PDB ID 5CHV, 5CHT) shed light on the role of ISG15 in USP18-STAT2 complex formation. It has been demonstrated that mUSP18 has an equilibrium of two distinct conformations, OPEN and CLOSED, which are characterized by distinct switching loop conformations (Fig. 3d)^28^. The OPEN conformation, with a flexible switching loop structure, allows mISG15 C-tail to access the mUSP18 catalytic site. In contrast, the switching loop in the CLOSED state adopts a short α-helix with mF132 positioned to obstruct the engagement of mISG15 to the mUSP18 catalytic site by generating steric hindrance with mR151/mR153 (Fig. 3d). The switching loop in the STAT2-USP18-ISG15 ternary complex cryo-EM structure resembles that of the mUSP18 OPEN state due to the presence of ISG15, causing side chains of F135, Q62, and Q137 of USP18 to orient toward P292/K295 of STAT2, promoting protein-protein interactions (Fig. 3e). The switching loop of USP18 could adopt OPEN or CLOSED conformations in the absence of ISG15. Based on the structure, it is speculated that the CLOSED state in which the Q62/F135/Q137 of USP18 shift away from the STAT2 interface will potentially reduce the affinity between USP18 and STAT2, leading to a less stable complex. This observation is supported by mutagenesis where F135E of USP18 showed reduced signal in the NanoBRET assay (Fig. 2g). The structural analysis and mutagenesis indicate that the OPEN conformation stabilized by ISG15 binding plays a significant role in the interactions between USP18 and STAT2. Overall, our ternary complex cryo-EM structure aligns with biochemical findings on USP18-STAT2, highlighting key amino acid residues and uncovering the mechanism by which ISG15 influences USP18-STAT2 interactions by modifying the conformational equilibrium.

### STAT2 negatively regulates USP18 catalytic function

The cryo-EM structure also revealed an unexpected conformational change within the USP18 catalytic triad residues that were not predicted by the AF2 model (Fig. 4a, Supplementary Fig. S4a). The active conformation of the catalytic triad (nucleophile-base-acid) is essential for cysteine protease catalysis including USP18. As shown in Fig. 4a, the active catalytic triad conformation revealed by the mUSP18-mISG15 crystal structure suggests the H-bond network and optimal geometry between mC61 (nucleophile), mH314 (base) and mN331 (acid) of mUSP18 are imperative^28^. Likewise, a similar conformation was also observed in the active USP30-UB complex structure (Fig. Supplementary Fig. S4b)^31^. Strikingly, K218 and R300 of STAT2 in the USP18-STAT2 complex are positioned near the catalytic triad residues of USP18, displacing the side chains of H318 and N335 from their active catalytic triad conformations (Fig. 4a, Supplementary Fig. S4c). This causes outward rotations of approximately 110° and 90° for H318 and N335, respectively (Fig. 4a), potentially making the USP18 catalytically inactive.

**Fig. 4.**
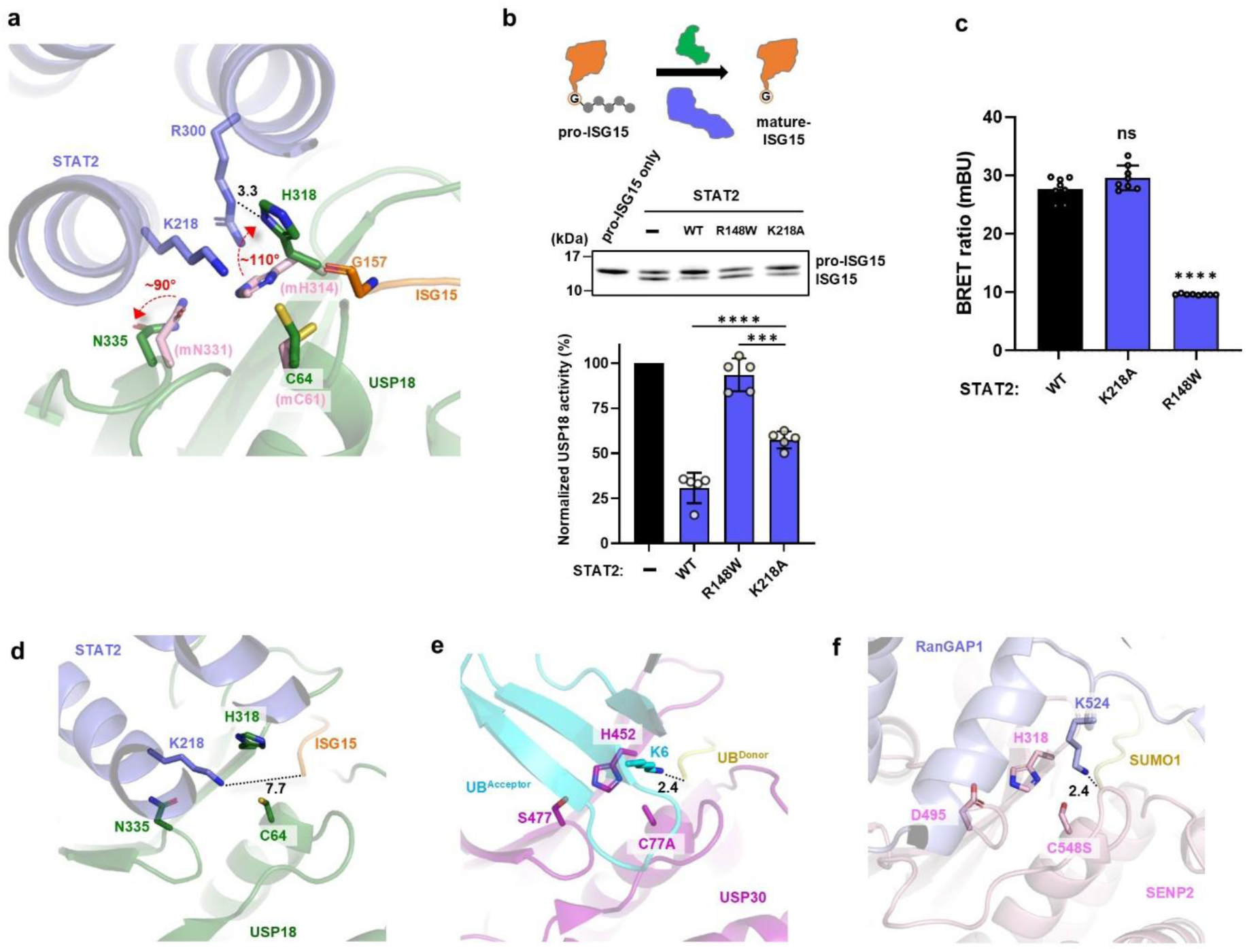
STAT2 inhibition for USP18 catalytic function. **a)** Catalytic triad in the cryo-EM structure (STAT2:blue, USP18:green, ISG15:orange) overlaying with the active catalytic triad nucleophile (mC61), base (mH314) and acid (mN331) from mUSP18-mISG15 crystal structure (PDB 5CHV) (mUSP18:pink). A hydrogen bond between USP18 H318 and STAT2 R300 is depicted by a black dash line. Rotation of His318 and N335 are noted with a red dash arrow. **b)** pro-ISG15 cleavage in the absence or presence of STAT2 variants. The scheme of assay is shown above. Data are mean ± s.d. (n = 5, biologically independent samples) with P-values obtained by ordinary one-way ANOVA test; *** and **** indicate significance at P = 0.0003 and < 0.0001 to STAT2 K218A, respectively. **c)** NanoBRET assay for STAT2 WT, R148W and K218A. Data are mean ± s.d. (n = 8, biologically independent samples) with P-values obtained by ordinary one-way ANOVA test; **** and ns indicate significance at P < 0.0001 and not significant, respectively. **d-f)** STAT2 K218 conformation (**d**) is compared with other cysteine protease and its substrate complex structures, USP30 (magenta) with K6-linked diUB (UB^Donor^:gold, UB^Acceptor^:cyan) (PDB 5OHP) (**e**) and SENP2(Pink)-SUMO1(Gold)-RanGAP1(Light blue) (PDB 2IO2) (**f**). Distances between target lysine and C-terminal glycine are denoted with a black dash line.

STAT2 R300 forms a hydrogen bond with USP18 H318 to stabilize the inactive conformation (Fig. 4a). Sequence alignment of STAT2 suggests that K218 is a highly conserved residue in vertebrate species, indicating K218 may play an important role in IFN-I signaling (Fig. Supplementary Fig. S4d). To assess the impact of STAT2 K218 on the catalytic function of USP18, we employed a pro-ISG15 biochemical cleavage assay. USP18 catalyzes removal of the eight additional amino acid residues from pro-ISG15 G157 to generate mature-ISG15^25,32^. The results showed that USP18 mediated cleavage was negatively regulated by the STAT2 WT protein (Fig. 4b). However, this inhibition was alleviated by the STAT2 R148W variant which likely has a compromised USP18 interactions (Fig. 4b). Intriguingly, the STAT2 K218A mutant partially restored the catalytic inhibition observed in the WT condition. In the NanoBRET assay, the K218A mutant displayed a NanoBRET signal comparable to that of STAT2 WT, suggesting that the partial functional rescue is likely attributed to the H318 conformational rearrangement allowed by the K218A mutation (Fig. 4c).

Next, we examined another hypothesis regarding the effect of STAT2 K218 on USP18 catalytic function, based on the observation that STAT2 K218, USP18 C64 and ISG15 G157 are close in the complex structure (∼8.0 Å) (Fig. 4d). This proximity suggests that STAT2 K218 may be a potential ISGylation site. We attempted to compare the ISGylation profile of STAT2 WT and K218A in HEK293 cells using a transient overexpression system but did not detect ISGylated STAT2 or deISGylation (data not shown). In contrast, comparing the STAT2-USP18-ISG15 complex structure with the USP30 and K6-linked diUB or SENP2-SUMO1-RanGAP1 complex structures that represent the active intermediate states of deubiquitination or deSUMOylation shows that the substrate Lys sidechains are located closer to the C-terminal Gly of UB or SUMO within 2.4 Å (Fig. 4e, 4f)^31,33^. In addition, both USP30 H452 (base)/ S477 (acid) and SENP2 H318 (base)/ D495(acid) form an active catalytic triad conformation whereas the ones in USP18 are likely inactive as discussed above. Finally, the projection of STAT2 K218 side chain toward the ISG15 C-terminal Gly is notably distinct from the ones in USP30 and in SENP2. While STAT2 K218 approaches ISG15 C-terminal Gly horizontally from the H318/N335 side, both acceptor UB K6 and RanGAP1 K524 make vertical contacts with their respective donor UB or SUMO1 C-terminal Gly from the C77/H452 or C548/H318 sides (Fig. 4d-f). While further experiments are needed, structural analysis suggests STAT2 K218 is likely not an ISGylation site.

### Distinct single point mutations to study USP18 catalytic/scaffolding functions

The cryo-EM structure and the biochemical characterizations elucidate the specific interactions between USP18 and STAT2, enabling further investigation into both scaffolding and catalytic functions within USP18 biology. In paticular, the profiles of the catalytically dead mutants of USP18, C64S and C64R, showed that USP18 C64R significantly affected STAT2 affinity, whereas C64S did not (Fig. 2g). Because USP18 I60N retains catalytic activity but alters scaffolding function, while C64S affects only enzymatic function, I60N and C64S mutants are classified as scaffolding-deficient and catalytic-deficient mutants, respectively. These two mutants, together with C64R, were further studied to assess their effects on ISG15, USP18, and STAT2 complex formation. Biolayer interferometry (BLI) results indicated that all USP18 mutants had comparable ISG15 binding affinity, despite an initial concern that the C64R mutation might prevent ISG15 C-terminal Gly tail engagement to the USP18 catalytic site (Fig. 5a, Supplementary Fig. S5a). Subsequently, these USP18 variants were tested using the USP18-STAT2 *in vitro* AlphaLISA assay to examine their effect on USP18-STAT2 interactions. The results showed substantial reduction of signal for the USP18 I60N and C64R mutants, consistent with the observations from the NanoBRET assay, while the C64S variant exhibited a signal comparable to WT (Fig. 2g, 2h, 5b). In the catalytic triad of USP18 shown in the cryo-EM structure, the modeled C64R variant does not appear to impact ISG15 C-tail engagement but is predicted to introduce a steric clash with USP18 H318 and STAT2 K218, which may contribute to the reduction of USP18-STAT2 interactions (Supplementary Fig. S5b). In contrast, the modeled USP18 I60N does not show any steric clash; however, the polar side chain of I60N may perturb the hydrophobic network on USP18, which plays a role in STAT2 interactions (Supplementary Fig. S5c). This is supported by the NanoBRET result observed with USP18 I60N and F135E mutants (Fig. 2g). To understand the effects of these point mutations in a cellular context, the downstream activation of IFN-I signaling was investigated by monitoring phosphorylation of STAT1 following the release of STAT2 from the negative regulation mediated by USP18. Consistent with the previous report, the catalytic-deficient mutant USP18 C64S did not increase pSTAT1 level, whereas the scaffolding-deficient mutant USP18 I60N upregulated IFN-I signaling as evidenced by the substantially increased pSTAT1 level (Fig. 5c)^25^. As expected, USP18 C64R also elevated pSTAT1 population, likely attributable to its decreased STAT2 binding affinity as suggested by the NanoBRET and AlphaLISA data (Fig. 2h, 5b). Intriguingly, neither C64R nor I60N reached the pSTAT1 level observed in USP18 KO (Fig. 5c), which may be attributable to the residual affinity as indicated by the AlphaLISA signal (Fig. 5b). Finally, these three distinct USP18 mutants were evaluated in a HAP1 cancer cell killing activity assay in the presence of IFN. Cell growth was strongly inhibited by USP18 KO, whereas the USP18 C64S mutation did not impact cell proliferation. In contrast, the USP18 I60N and C64R mutations partially inhibited cell growth, phenocopying the USP18 KO, and are consistent with findings from the pSTAT1 assay (Fig. 5d). These profiles in phosphorylation of STAT1 and cancer cell killing suggested that the C64R phenotype is similar to I60N but it is catalytically inactive. Together, these results allowed us to clarify the roles of USP18 C64S, I60N and C64R mutants as tools to further explore USP18’s catalytic and scaffolding functions in IFN-I signaling. In sum, we propose that C64S functions as a catalytic-deficient mutant, I60N represents a scaffolding-deficient mutant, and C64R exhibits deficiencies in both catalytic and scaffolding activities to allow studying of the combinatory effect.

**Fig. 5.**
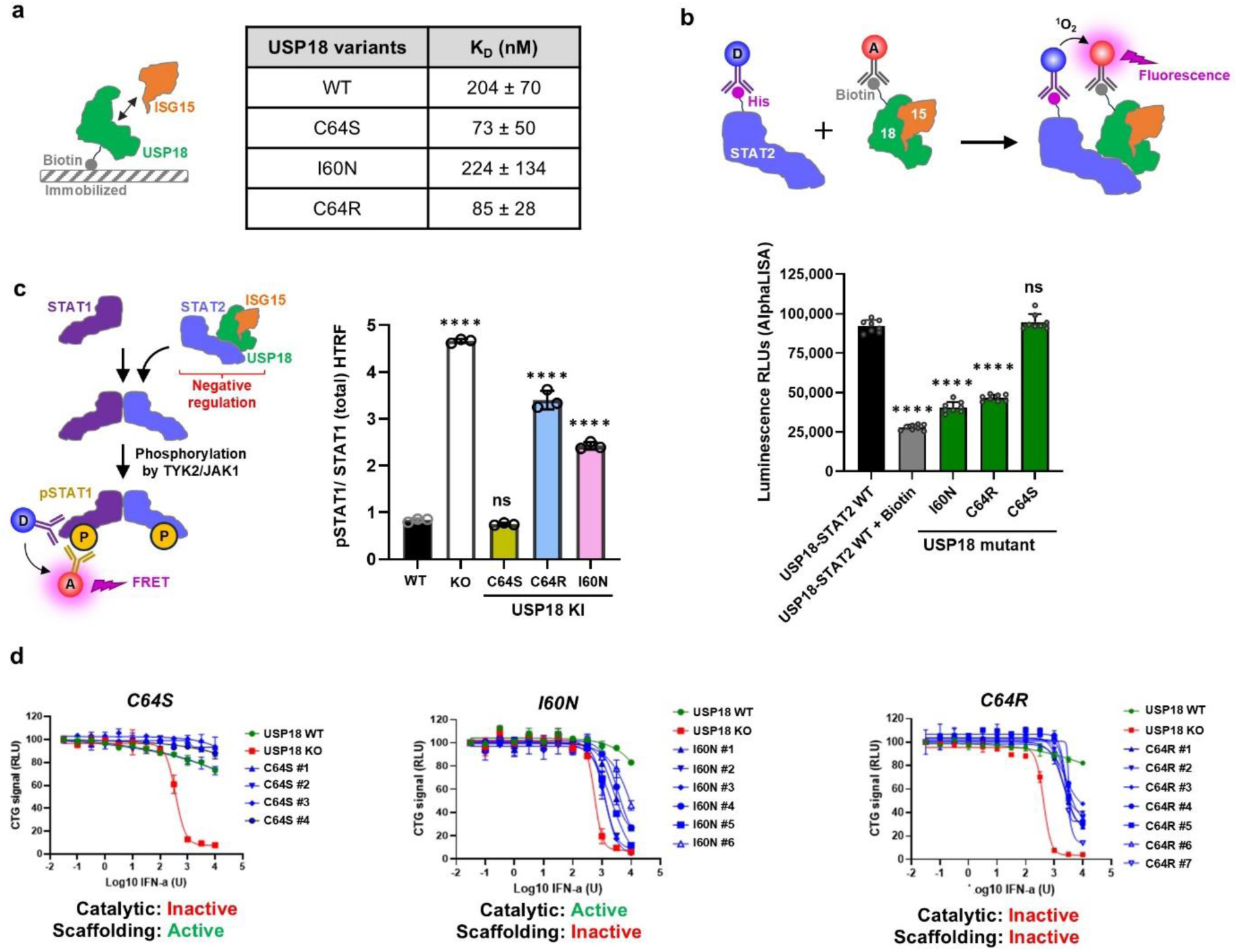
Distinct USP18 point mutants to explore USP18 catalytic and/or scaffolding functions. **a)** USP18-ISG15 BLI for USP18 variants. Data are mean ± s.d. (n = 2, biologically independent samples). The assay scheme is shown on left. **b)** AlphaLISA of USP18 variants for USP18-STAT2 interactions. Data are mean ± s.d. (n = 8, biologically independent samples) with P-values obtained by ordinary one-way ANOVA test; **** and ns indicate significance at P < 0.0001 and not significant to USP18-STAT2 WT, respectively. The assay scheme is shown above. **c)** Monitoring IFN-induced phosphorylation of STAT1 (pSTAT1) over total STAT1 level in HAP1 cells using parent (WT), USP18 KO, USP18 I60N, C64S, and C64R knock-in (KI) clones. Data are mean ± s.d. (n = 3, biologically independent samples) with P-values obtained by ordinary one-way ANOVA test; **** and ns indicate significance at P < 0.0001 and not significant to WT, respectively. The assay scheme is shown on left. **d)** Viability of USP18 C64S KI (left), USP18 I60N KI (middle) or USP18 C64R KI (right) HAP1 cells at the indicated IFN-α concentrations. The status of USP18 catalytic or scaffolding functions are denoted below.

## Discussion

USP18-STAT2 plays a pivotal role in the negative regulation of the IFN-I signaling, however, a detailed molecular understanding had not been fully uncovered due to the lack of structural information, particularly regarding the STAT2-USP18-ISG15 ternary complex. In this study, we present the unprecedented ternary complex structure of STAT2-USP18-ISG15 determined by cryo-EM at 3.05 Å resolution, facilitated by AlphaFold AI prediction and protein engineering via SPY-tag technology (Fig. 1). Complexes with transient or weak protein-protein interactions have been challenging for structural determination. To address this, the SPY-tag, known by its small size and rapid irreversible isopeptide bond formation, was leveraged to stabilize the USP18-STAT2 complex^26,27^. AlphaFold predictions are a valuable resource, but their models require cautious interpretation because predictions for side chain conformation may not be reliable for transient interactions or dynamic environments. As demonstrated here, AlphaFold2 successfully predicted the relative orientation of STAT2-USP18-ISG15. However, the AI-based prediction did not capture accurate side chain rearrangement, as evidenced by the large deviation in the catalytic triad conformation compared to that observed in the cryo-EM structure (Supplementary Fig. S4a).

Our structural-based mutagenesis provided the first evidence that the USP18-STAT2 complex negatively regulates the catalytic function of USP18, which may contribute to further shifting USP18 functions from enzyme to scaffolding (Fig. 4a-c). The co-localization of STAT2 K218 with the catalytic triad of USP18 presents the possibility that STAT2 K218 might be an ISGylation site for USP18. However, the significant structural deviation from the USP30-diUB and SENP2-SUMO1-RanGAP1 complexes which revealed how acceptor UB or substrate RanGAP1 Lys side chains engage the active site for donor UB or SUMO1 cleavage and transfer, indicates that STAT2-USP18-ISG15 may not adopt an active conformation^31,33^ (Fig. 4d-f). In addition, due to the large distance observed between ISG15 C-terminus and K218 of STAT2, the likelihood for ISGylation at STAT2 K218 is low, though further experiments are needed to address this question. Intriguingly, according to the PhosphoSitePlus database, STAT2 K218 has been identified as a ubiquitination site (Supplementary Fig. S4e)^34,35^. If STAT2 K218 serves as a ubiquitination site, the impact of ubiquitination on STAT2 K218 on USP18 function regulation remains to be determined^34,35^. One plausible hypothesis suggested by the cryo-EM structure is that if UB molecules are conjugated to STAT2 at residue K218, significant steric clash would occur with USP18 or ISG15, thereby preventing USP18 from interacting with STAT2, which may lead to upregulation of pSTAT1 and cell death.

Active site arginine mutations have been shown effective for inactivating DUB enzymes to study their functions^36^. In UBP8 and OTUD1, arginine mutations at the catalytic cysteine reduced affinity toward UB or UB chains, likely due to steric clashes with the UB C-terminal glycine tail or the isopeptide bond between donor and acceptor UB molecules introduced by the arginine replacement^36^. In the case of USP18, C64R mutations did not diminish its affinity for ISG15; however, it markedly affected interactions with STAT2. Structural analysis suggests that the substituted arginine side chain may cause steric hindrance with both the USP18 catalytic triad residue H318 and STAT2 K218 (Fig. 5a, 5b, Supplementary Fig. S5b). The C64R structural model and the C64R phenotype comparison with I60N and C64S allowed us to classify them into catalytic/scaffolding dual-deficient, scaffolding-deficient, and catalytic-deficient mutants (Fig. 5c, d). These characterizations provided valuable tools to investigate USP18’s DUB activity and IFN sensitivity. The USP18 C64R mutant has previously been characterized as catalytically inactive due to its lack of activity in an ISG15 activity-based probe assay^37^. In this report, USP18 C64R showed cancer cell killing activity upon IFN treatment, different from the other catalytic-deficient mutant C64S. Our cryo-EM and mutagenesis findings suggest that in addition to its impact on USP18 catalytic function, C64R mutation introduces a steric clash with STAT2 (Supplementary Fig. S5b), leading to a loss of scaffolding function as the underlying mechanisms to the observed increased IFN sensitivity, consistent with the other scaffolding-deficient mutations described in this study (Fig. 5).

DUBs are associated with several human diseases, including cancer, neurodegenerative, and autoimmune diseases^38^. Despite their significance, developing drugs that target DUBs has remained challenging for decades because of complex enzyme regulation and selectivity issues. However, as exemplified by USP7, USP14, and UCHL1, the significant learnings in DUB biology and technological advancements have enabled identification of small molecule inhibitors which modulate DUB functions^38–42^. As suggested in this study, loss of USP18-ISG15 interactions attenuates USP18-STAT2 affinity (Fig. 3) affecting the negative regulation of IFN-I signaling by USP18 scaffolding function. Finding a chemical matter that displaces ISG15 from USP18 would be a promising pathway for a cancer therapy targeting USP18. An alternative intriguing approach is to seek a covalent ligand targeting a reactive amino acid on the protein surface such as a cysteine residue^43,44^. Covalent ligands are often found through either structure-based drug design (SBDD) or covalent chemical library screening efforts. In the case of OTUB1, a K48-linked ubiquitin-specific DUB, EN523 was identified via cysteine-reactive library screening^45^. This approach is utilized in the development of deubiquitinase-targeting chimeras (DUBTACs) to stabilize target protein levels by promoting proximity-induced deubiquitination^45^. As shown in Fig. 2f and 2h, USP18 has several surface cysteine residues that participate in USP18-STAT2 interactions, and mutations at these positions significantly reduced complex formation. Although further studies are needed, these surface cysteine residues, including the catalytic Cys64, may represent a potential target for covalent drugs aimed at disrupting USP18-STAT2 interactions to unleash the IFN response. Overall, the cryo-EM structure and biochemical characterization of the STAT2-USP18-ISG15 ternary complex described herein clarify how USP18’s scaffolding and catalytic activities are regulated by its interactions with STAT2 and ISG15. These insight into USP18’s role in negative regulation of IFN-I signaling provide multiple avenues to target USP18 for potential cancer therapy.

## Materials and Methods

### Protein expression and purification

USP18(16-372) (Q9UMW8) and ISG15(1-157)C78S (P05161) (refer ISG15 in this study) were purified as described previously^25^. STAT2(130-706) (P52360) wild-type and mutants were inserted into pET28a(+) vector (EMD Millipore 69864-3) with N-terminal 6xHis-tag and Tobacco Etch Virus protease cleavage site and C-terminal Avi-tag. For cryo-EM, N-terminal 6xHis-SpyCatcher-STAT2(1-706), 6xHis-SpyTag-USP18(26-372) were inserted into pFastbac vector. STAT2(130-706) variants were expressed in Escherichia coli BL21(DE3) cells and purified by Ni-affinity chromatography followed by size exclusion chromatography (SEC) with a SEC buffer (25mM HEPES pH 8.0, 200mM NaCl, 1.0mM TCEP). SPY-tagged STAT2-USP18-ISG15 complex were co-expressed in Spodoptera frugiperda (Sf9, ATCC CRL-1711) insect cells and purified by tandem affinity purification using Ni-beads (STAT2-USP18) then GST-beads (ISG15). After the GST-tag cleavage by Prescission protease, the complex was polished by SEC. For AlphaLISA assay, USP18(16-372) variants with N-terminal 6xHis and C-terminal Avi tag was designed and co-purified with GST-ISG15. The Avi-tag is fully biotinylated by co-expressing with a biotin ligase (BirA) in insect cells. After the tandem purification using Ni- and GST-beads followed by 6xHis and GST tag removal by Prescission protease, the biotiylated Avi-tagged USP18-ISG15 complex was further purified by SEC. For BLI, the biotiylated Avi-tagged USP18 variants and untagged ISG15 were purified as shown previously^25^.

### Mass photometry

Mass photometry measurements were done with the TwoMP mass photometer (Refeyn), calibrated as per manufacturer protocols. All measurements were performed with a total volume of 20 μl in the measuring well: adding 10 μl of SEC buffer to clean wells before adding a protein sample to make a final concentration of 100 nM. Movies of scattering were recorded using AcquireMP (version 2023 R2.2) and were subsequently analyzed by DiscoverMP (version 2023 R2). The optical contrast value at the particle center was compared to the calibration curve obtained from a control sample comprising 65 and 130 kDa proteins to determine the mass of particles.

### Cryo-EM

#### Sample vitrification and data collection

The STAT2-USP18-ISG15 complex with the SpyCatcher tag was diluted to 0.80 mg/mL with SEC buffer. The purified complex was applied onto a glow discharged Quantifoil R1.2/1.3 200 mesh Au grids and flash frozen into liquid ethane using the Vitrobot Mark IV (ThermoFisher Scientific). The grids were loaded into the Titan Krios G2 transmission electron microscope equipped with a Falcon 4i direct detector and Selectris energy filter (ThermoFisher Scientific). About 16,500 movies were acquired using EPU (version 3.6.0.6389) in counting mode with AFIS, a pixel size of 0.75 Å/pixel, 10 eV slit and a total dose of 40 e-/Å^2^ over a defocus range of −0.8μm to −2.8μm with 2.81 second exposure.

#### Data processing

All data processing procedures were done using CryoSPARC v4.6.0^46^ and shown in Supplementary Fig. S2a. The movies were patch motion corrected to generate averaged micrographs, where were used to patch CTF estimation. Particles were initially picked using blob picker and classified into 2D class averages for Topaz template auto-picking^47^. After the first round of classification, five class averages were selected for Topaz training and Topaz auto-picking resulting in about 2.1 million particles picked. These particles were extracted with a box size of 360 pixels and were sorted through multiple rounds of 2D classifications providing about 527,700 particles for 3D reconstruction, 3D classification and refinement. *Ab-initio* reconstruction generated four initial models with one model resembling a complex while the other three are noise. A 3D homogeneous refinement was done revealing a low resolution structure that can fit the mUSP18-mISG15^28^ and hSTAT2^48^ structures as a starting point for modeling. Since the *Ab-initio* reconstruction revealed noisy particles in the stack, the particles underwent 3D classification to further sort out high-resolution and low-resolution particles. One of the three 3D classes containing 143,496 particles provided a reconstruction with domains that are best resolved. The particles from that 3D class underwent multiple rounds of homogenous refinement, non-uniform 3D refinement and reference-based motion correction refining a final STAT2-USP18-ISG15 structure to 3.05Å.

#### Atomic modeling and model refinement

The structures for mUSP18-mISG15 (PDB: 5CHV)^28^ and hSTAT2 (PDB: 6WCZ)^48^ were fitted into the final cryo-EM structure using UCSF Chimera^49^. The atomic model was built with minor refinements in COOT^50^ using the fitted structure as a guide. Phenix real space refinement was performed after modeling was completed to further optimize the model. The final refinement statistics are listed in Supplementary Table 1. The structural comparison and figure generation in this manuscript were done with the graphic program PyMOL^51^. Solvent accessible surface area was calculated using AREAIMOL in CCP4 suite^52^.

#### Nano-Luciferase Bioluminescence Resonance Energy Transfer (NanoBRET)

Protocol is based on the NanoBRET Protein-Protein Interaction System Technical Manual published by Promega (Cat. #N1661). Hek293T cells were cultured using standard cell culture techniques and 400,000 cells were plated into each well of a 12-well plate. After 4 hours, cells were transfected using PEI (Polysciences #239661) pH 7.0 at 1 mg/ml. For each transfected well, 3 μg of PEI was diluted in 62 μl of OptiMEM I Reduced Serum Medium, with phenol red. In another 62 μl of OptiMEM, 0.1 μg of USP18 and STAT2 constructs and 1 μg of ISG15 construct was added. The PEI mixture was then added to the DNA mixture, mixed by pipetting, and incubated at room temperature for 20 min. The DNA/PEI complexes were added dropwise to the appropriate well and the plate was rocked to mix. Cells were incubated at 37 °C for 24 hours.

After 24 hours, media was removed from cells and media containing the HaloTag NanoBRET 618 ligand was added (Ligand Media: Opti-MEM I Reduced Serum Medium, no phenol red (Thermo #11058-021)+ 4% FBS at a 1:1000 ratio of 618 ligand/media, Promega N1661). Cells incubated at 37 C° overnight. The next day, media was removed and 200 μl of TrypLE with no phenol red (Thermo #12604-013) was used to lift cells. Cells were diluted to 200,000 viable cells/mL in Ligand Media and 25uL was plated into each well of a 384 well plate (Greiner 78108). Cells were spun down at 1000xg for 30 seconds. A 5x substrate solution was prepared by diluting NanoBRET Nano-Glo substrate (Promega N1661) by 100x in Opti-MEM I Reduced Serum Medium, no phenol red. 6 μl of this substrate solution was added to each well of the 384 well plate, plates were centrifuged at 1000xg, and donor (460nm) and acceptor (618nm) emission was measured using an Envision 2104 Multilabel Plate Reader (Mirror: Luminescence 404; Emission filters: 600LP NanoBRET, M460/50 nm NanoBRET; Measurement time: 0.1 second). NanoBRET ratio was calculated by dividing the acceptor emission by the donor emission for each well. Mean ± s.d. with P-values were obtained using GraphPad Prism (version 10.4.1).

#### Biolayer Interferometry (BLI)

Binding assays for USP18 and ISG15 complex formation were performed on an Octet Red384 biolayer interferometry instrument (ForteBio, Inc.) as previously described ^25^ with minor modifications. In brief, biotinylated Avi-tagged USP18 proteins immobilized on streptavidin-coated sensors were put into wells containing ISG15 (0.14 to 4 mM in a 33% dilution series) for a 5-minute association, followed by a 5-minute dissociation in ISG-free wells. Rates of association and dissociation were determined using GraphPad Prism (version 10.4.1) and the average of two determinations each were used to calculate K_D_ values.

#### Pro-ISG15 cleavage assays

The assays were performed as previously reported with minor modifications^25^. 1 μM recombinant USP18 and 5 μM recombinant pro-ISG15 were incubated for 15 minutes at 37 °C in the presence or absence of 5 μM STAT2 (WT, R148W, K218A). Alternation from pro-ISG15(1-165) to mature ISG15(1-157) was monitored by SDS-PAGE and normalized USP18 activity was calculated by quantifying band intensities using Image J (version 1.47v). Mean ± s.d. with P-values were obtained using GraphPad Prism (version 10.4.1).

#### AlphaLISA

Biotinylated Avi-tagged USP18(16-372)/ISG15(1-157) complex were used at equimolar concentration of 30 nM and His-tagged STAT2(130-706) was utilized at 300 nM concentration. The samples were diluted by a buffer of 20 mM Tris-HCl, pH 7.5 150 mM NaCl 0.05 % CHAPS With 0.5 mM TCEP. These were mixed with 20 μg/mL AlphaLISA donor (Anti-6xHis Alpha Donor Beads supplier part number AS116D) and acceptor (Streptavidin AlphaLISA Acceptor Beads supplier part number AL125C) beads, 20 μl of the complete mix was plated in 3824 Costar 384 well plates and incubated for 60 minutes before being read on an Envision reader for AlphaLISA signal at Excitation Wavelength Laser (nM) 680 and Emission Wavelength 575 with excitation time 180 ms and detection time 370 ms. An excess amount of biotin was added to saturate the acceptor beads to prevent it from binding STAT2. Mean ± s.d. with P-values were obtained using GraphPad Prism (version 10.4.1).

#### Cellular pSTAT1/total STAT1 Homogeneous Time Resolved Fluorescence (HTRF) assay

The assays were performed as previously reported with minor modifications^25^. pSTAT1 and total STAT1 levels in USP18 KO, I60N KI, C64S KI or C64R KI clones were analyzed and the ratio of pSTAT1/total STAT1 levels were calculated accordingly. Mean ± s.d. with P-values were obtained using GraphPad Prism (version 10.4.1).

#### CellTiter-Glo assay

Hap1 USP18^C64S^, USP18^I60N,^ and USP18^C64R^ cells were subjected to an 11-point dose response of Human IFN-Alpha2b (pbl Assay Science, 11105-1) with a top concentration of 9,000 U/mL and 3.33 dilution series. Cells were plated into 384-well white microplates (Corning, 3570) at 3,000 cells/well in 50 µl media containing Human IFN-Alpha2b and incubated for 72 hours. at 37 °C/ 6% CO_2_. Cells were lysed with 50 µl CellTiter-Glo 2.0 (Promega, G9243) and incubated in the dark for 20 minutes at room temperature. Luminescence was measured on the GloMax Discover (Promega, GM3000). Generation of these genetically modified cell lines have been described previously^25^.

## Data availability

The sequence of WT human USP18, ISG15 and STAT2 proteins are available in UniProt with the accession code Q9UMW8, P05161 and P52630 respectively. The final STAT2-USP18-ISG15 complex cryo-EM map and model are deposited in the Electron Microscopy Data Bank (EMDB) under accession code EMD-74146 and Protein Data Bank (PDB) under accession code 9ZFO. Additionally, data from this finding is available upon request from the corresponding author.

## Acknowledgements

We appreciate Adan Pinto Fernandez and Benedikt Kessler for discussion, and Amir Arellano-Saab for cellular ISGylation experiment supports.

## Contributions

K.W.H., R.P., D.R.H., V.J., E.C.R., C.L., Heather W., K.L., Z.H., K.F., R.A.C., and M.Y. performed experiments and data analysis. T.C., P.D.W., P.M.L., M.C., S.H., Huixian W., F.W. and M.Y. supervised experiments and data analysis. S.H. supervised cryo-EM laboratory operation. P.D.W., F.W. and M.Y. conceived, designed and supervised the project. K.W.H., Huixian W. and M.Y. wrote the manuscript. All authors analyzed and discussed the results and approved the manuscript.

---

**Supplementary Fig. S1.**
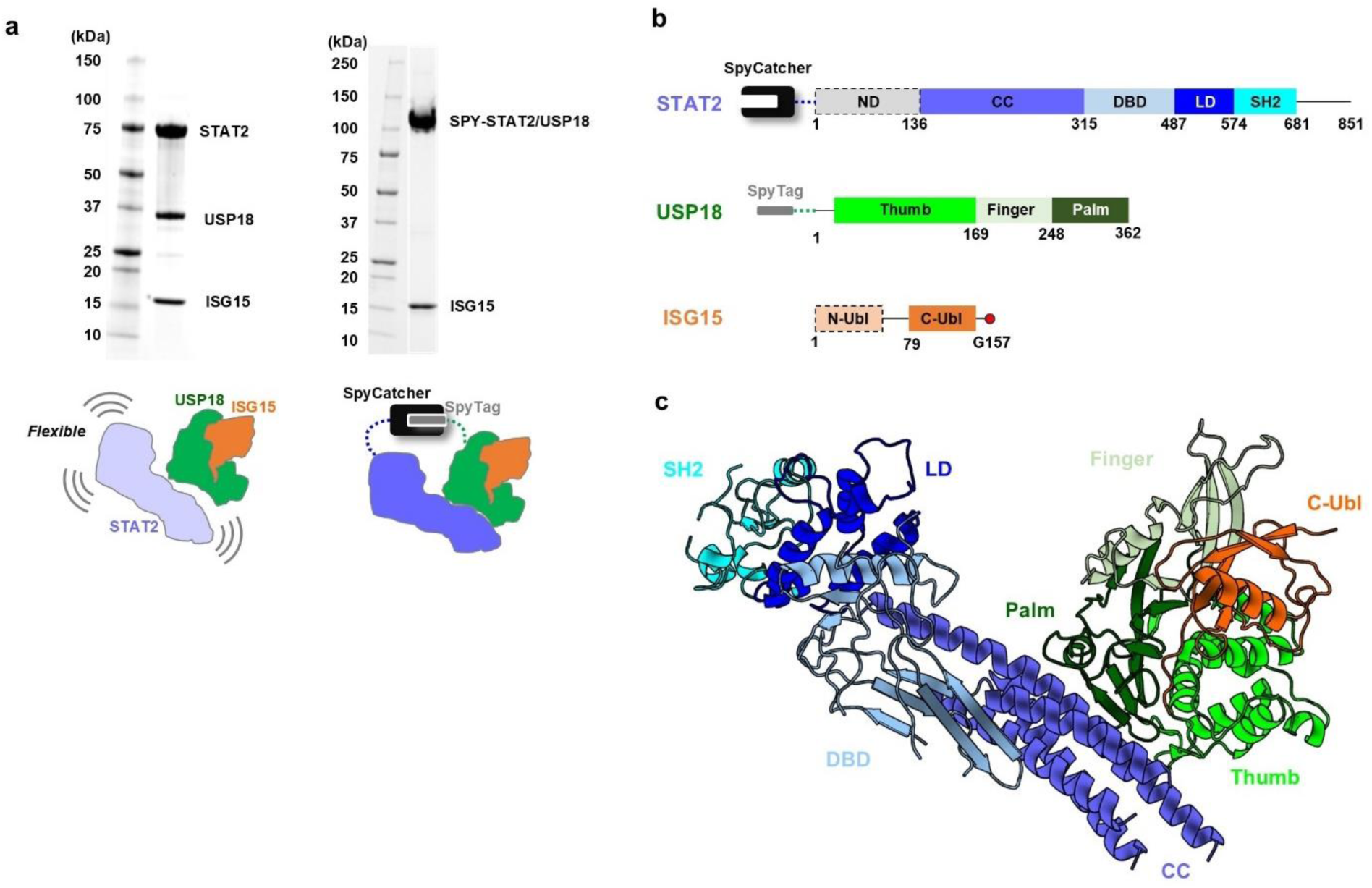
Preparation of cryo-EM samples. **a)** Comparison of WT and SPY-complex cryo-EM samples. SDS-PAGE gels represent the MW shift of USP18-STAT2 due to the covalent linkage of SpyCatcher/SpyTag. **b)** Schematic view of SPY-complex cryo-EM sample design with illustrating functional domains for STAT2, USP18 and ISG15. STAT2: ND, N-terminal domain; CC, coiled-coil domain; DBD, DNA-binding domain; LD, linker domain; SH2, Src Homology 2 domain. USP18: Thumb, Palm and Finger regions are canonical classification for deubiquitinase (DUB). ISG15: two Ubiquitin-like domains at N- or C-terminus (N-Ubl, C-Ubl). SpyCatcher/SpyTag are fused at N-terminus of STAT2 and USP18 respectively. **c)** Functional domains mapped on the STAT2-USP18-ISG15 complex structure. The color scheme is same as in Supplementary Fig. S1b.

**Supplementary Fig. S2.**
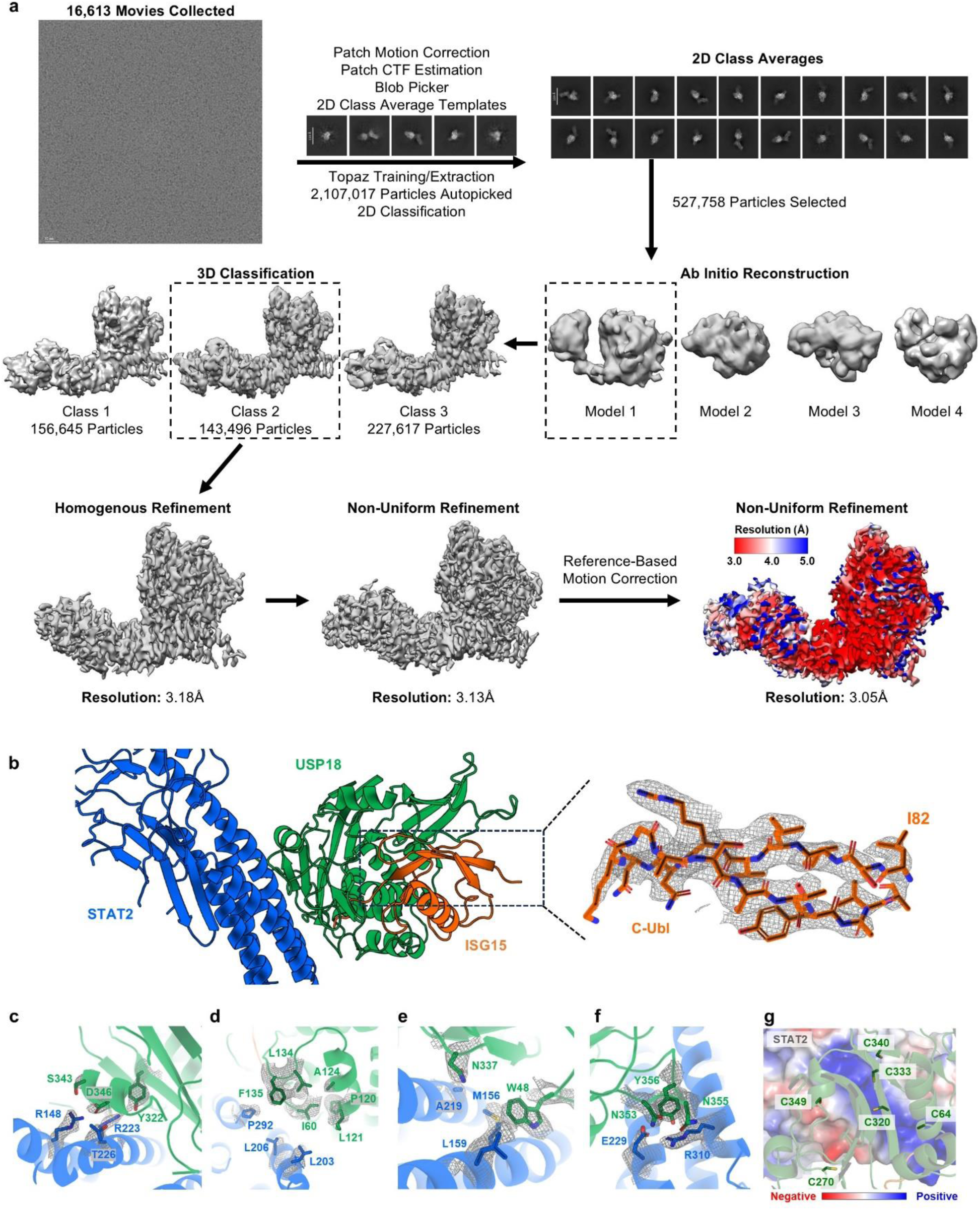
Cryo-EM workflow and models. **a)** Flow chart of cryo-EM data processing including a representative micrograph, representative 2D class averages and 3D reconstructions showing distinct views and features of the complex. The final cryo-EM map is colored by local resolution. **b)** Model fitting into the cryo-EM map highlighting ISG15 N-Ubl domain is missing. **c-f)** Corresponding cryo-EM maps for residues in mutagenesis study in Fig. 2. **g)** Cysteine patch of USP18 with showing the electrostatic map of STAT2.

**Supplementary Fig. S3.**
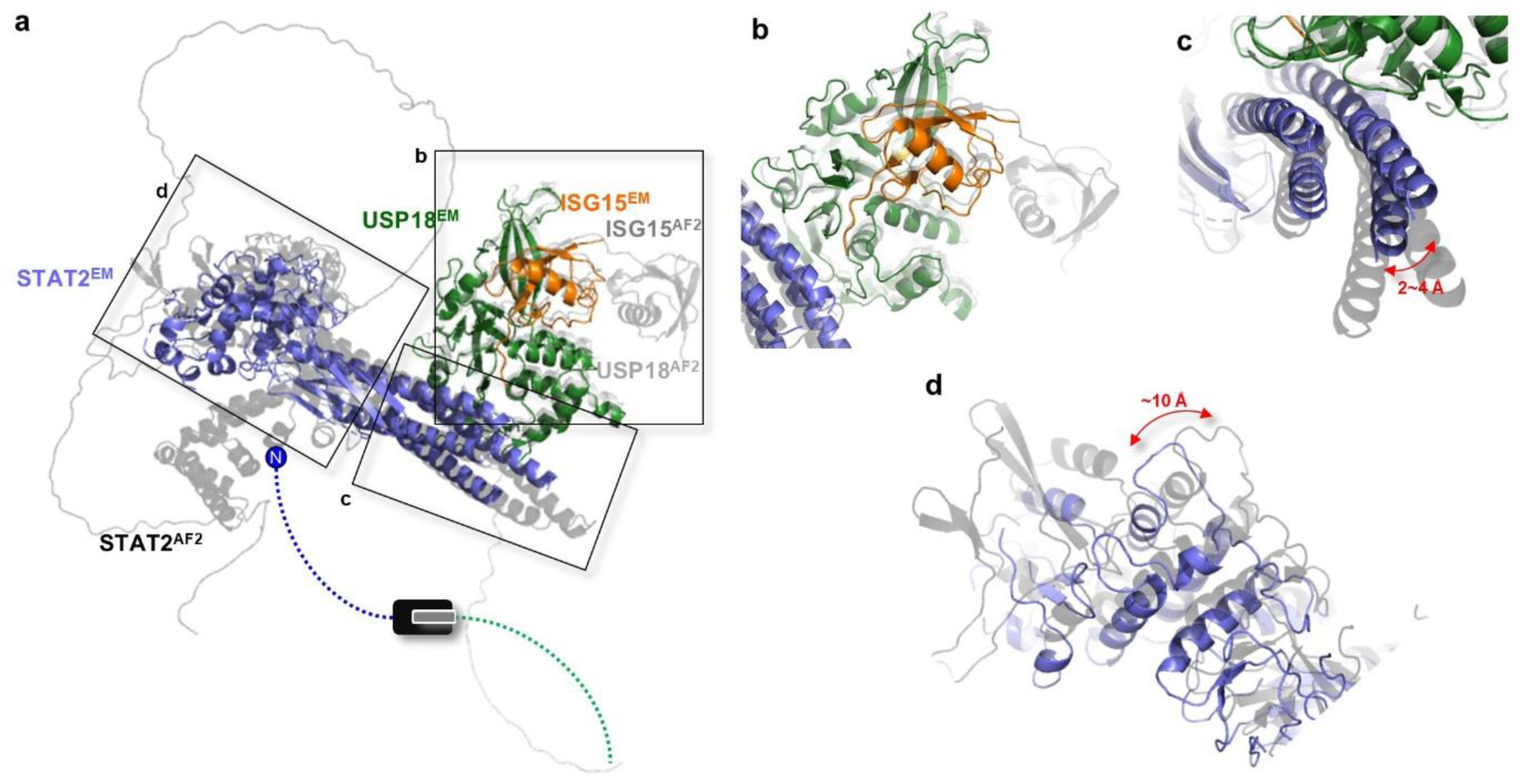
Cryo-EM structure and AF2 model comparison. **a)** The AF2 model (STAT2:gray, USP18:black, ISG15:light gray) was superimposed onto the cryo-EM structure (STAT2:blue, USP18:green, ISG15:orange). **b)** USP18-ISG15 has marginal variation but lacks ISG15 N-Ubl. **c)** STAT2 CC helices tilted toward USP18. **d)** STAT2 LD/SH2 exhibited a large conformational shift.

**Supplementary Fig. S4.**
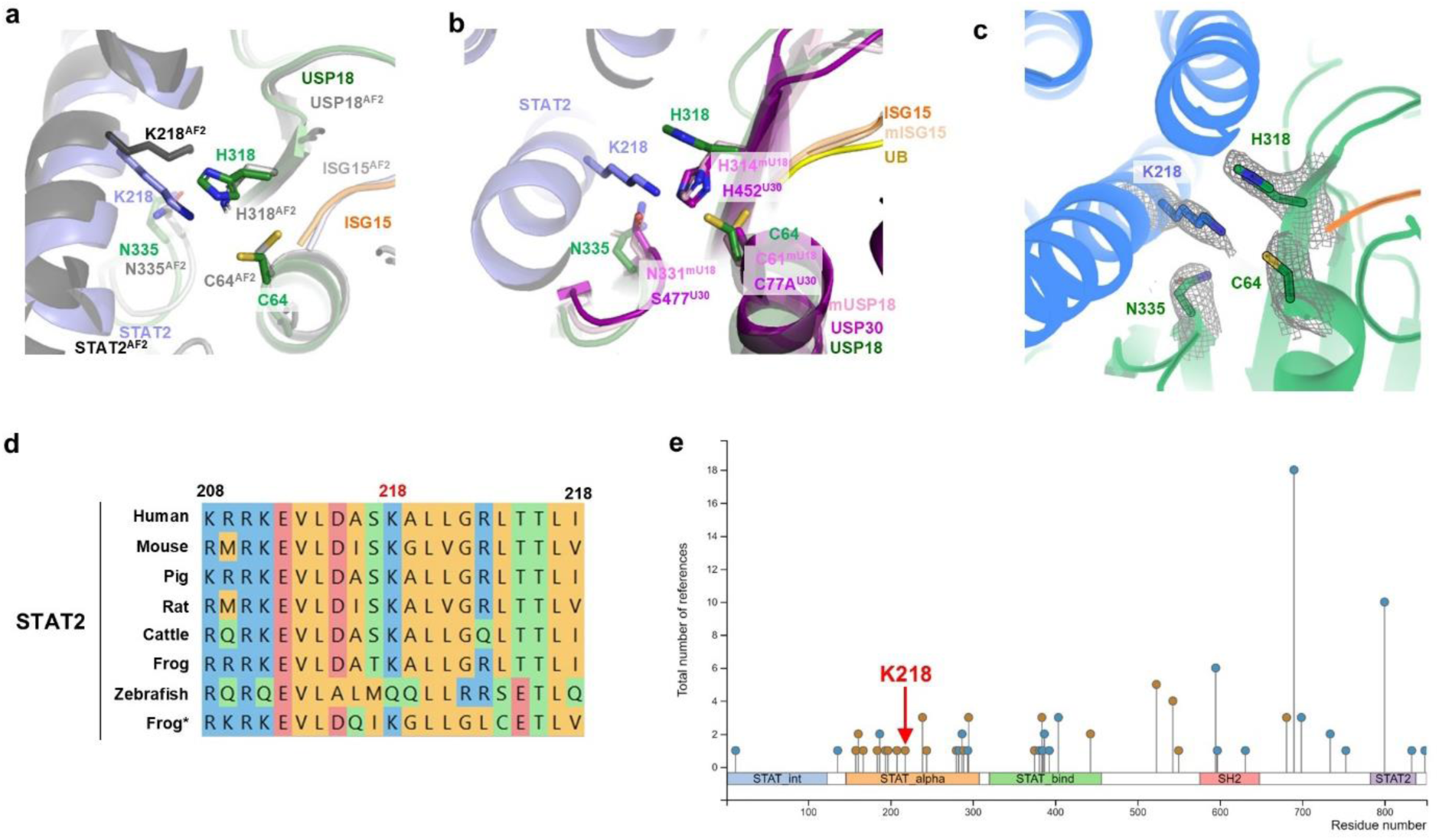
Insight into the functional role of STAT2 K218. **a)** The catalytic triad conformations in the cryo-EM structure with the AF2 model overlayed. Color scheme is same as in Supplementary Fig. S3. **b)** The catalytic triad conformation in the cryo-EM structure was compared with the active catalytic triad conformations in mUSP18-mUSP15 (PDB 5CHV) and USP30-UB (PDB 5OHP). **c)** The cryo-EM map supporting the STAT2 K218 and USP18 H318 conformations. **d)** STAT2 sequence alignment among several vertebrates highlighting K218 is conserved well. **e)** Post-translational modifications of STAT2 in PhosphositePlus database^34^. Ubiquitination, phosphorylation or acetylation sites are colored orange, blue or green respectively.

**Supplementary Fig. S5.**
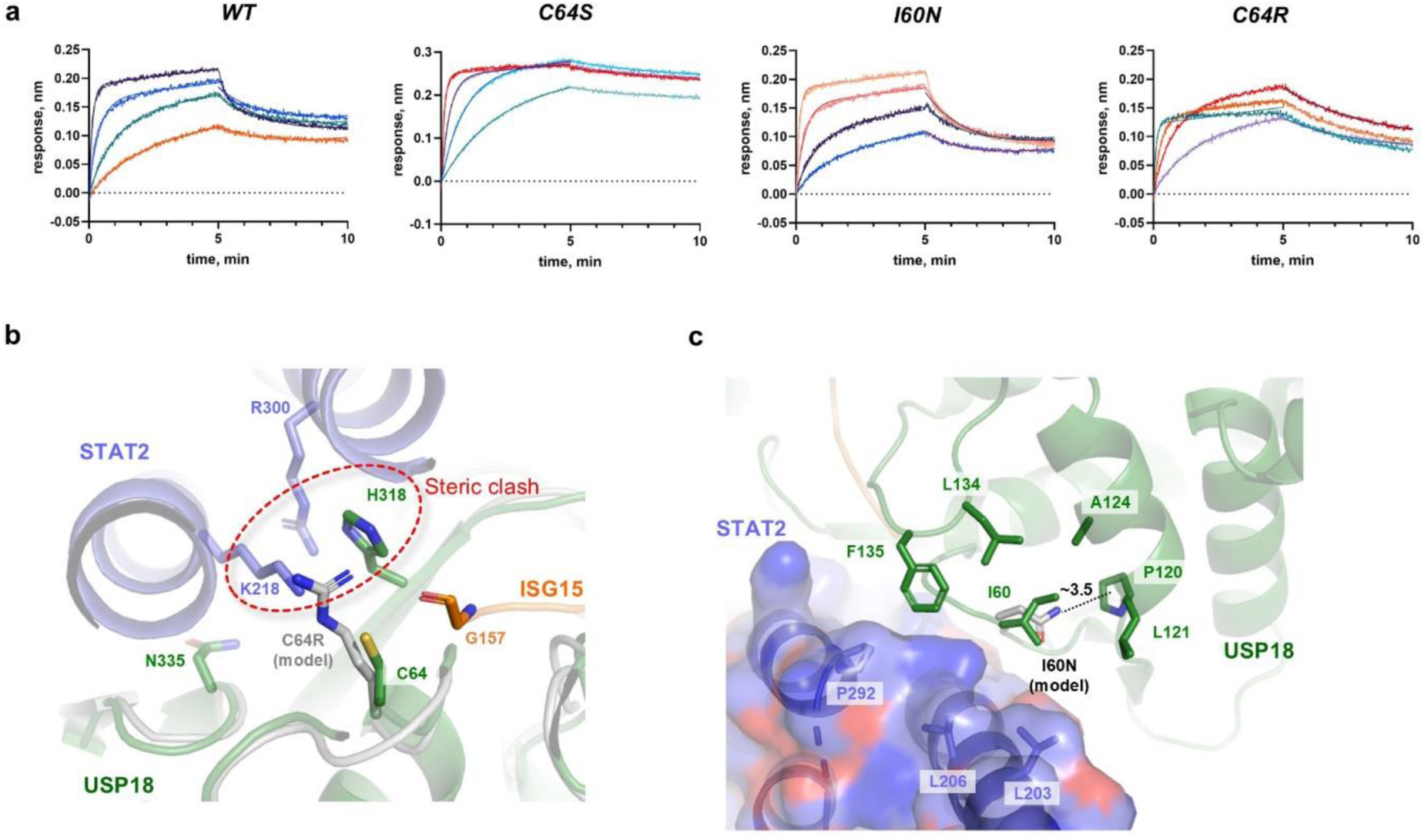
Characterization of USP18 C64R and I60N. **a)** The representative BLI sensorgram analyzed in Fig. 5a. **b)** The model of USP18 C64R showing a steric clash with USP18 H318 and STAT2 K218. **c)** The potential perturbation in the USP18 I60 hydrophobic network caused by I60N mutation.

**Supplementary Table 1.** Cryo-EM data collection, refinement and validation statistics.

|  | STAT2-USP18-ISG15<br>(EMD-74146)<br>(PDB 9ZFO) |
| --- | --- |
| **Data collection and processing** |  |
| Magnification | 165,000 |
| Voltage (kV) | 300 |
| Electron exposure (e-/Å<sup>2</sup>) | 40.0 |
| Defocus range (µm) | -0.8 to -2.4 |
| Pixel size (Å) | 0.750 |
| Movies recorded | 16,613 |
| Symmetry imposed | C1 |
| Initial particle images (no.) | 2,107,017 |
| Final particle images (no.) | 143,496 |
| Map resolution (Å) | 3.05 |
| FSC threshold 0.143 |  |
| Map resolution range (Å) | 3.0-3.7 |
| **Refinement** |  |
| Initial model used | PDBs: 5CHV and 6WCZ |
| Model resolution (Å) | 3.05 |
| FSC threshold |  |
| Model resolution range (Å) | N/A |
| Map sharpening *B* factor (Å<sup>2</sup>) | -116.15 |
| Model composition |  |
| Non-hydrogen atoms | 6.635 |
| Protein residues | 827 |
| Ligand | 0 |
| *B* factors (Å<sup>2</sup>) |  |
| Protein | 73.32 |
| Ligand | N.A. |
| R.m.s. deviations |  |
| Bond lengths (Å) | 0.005 |
| Bond angles (°) | 0.954 |
| Validation |  |
| MolProbity score | 1.60 |
| Clashscore | 4.35 |
| Poor rotamers (%) | 0.0 |
| Ramachandran plot |  |
| Favored (%) | 94.40 |
| Allowed (%) | 5.60 |
| Disallowed (%) | 0 |

